# Transcranial stimulation enhances memory-relevant sleep oscillations and their functional coupling in mild cognitive impairment

**DOI:** 10.1101/095588

**Authors:** Julia Ladenbauer, Josef Ladenbauer, Nadine Külzow, Rebecca de Boor, Elena Avramova, Ulrike Grittner, Agnes Flöel

**Author notes:** These authors contributed equally to this work. Correspondence: Julia Ladenbauer or Agnes Flöel, Charité Universitätsmedizin Berlin, Charitéplatz 1, 10117 Berlin, Germany.

## Abstract

Alzheimer’s disease (AD) not only involves loss of memory functions but also prominent deterioration of sleep physiology, already evident in the stage of *mild cognitive impairment* (MCI). Cortical slow oscillations (SO, 0.5-1 Hz) and thalamo-cortical spindle activity (12-15 Hz) during sleep, and their temporal coordination, are considered critical for memory formation. We investigated the potential of slow oscillatory transcranial direct current stimulation (so-tDCS), applied during a daytime nap in a sleep state-dependent manner, to modulate these activity patterns and sleep-related memory consolidation in 16 patients with MCI.

Stimulation significantly increased overall SO and spindle power, amplified spindle power during SO up-phases, and led to stronger synchronization between SO and spindle power fluctuations in electroencephalographic recordings. Moreover, visual declarative memory was improved by so-tDCS compared to sham stimulation, associated with stronger synchronization. These findings indicate a well-tolerated therapeutic approach for disordered sleep physiology and deficits in memory consolidation in MCI patients.

## Introduction

Difficulties in forming and retrieving episodic memories are noted early in the course of Alzheimer’s disease (AD) and constitute a core component of the condition (Bäckmann et al., 2004). Likewise, sleep disturbances appear to be a major characteristic of AD-related dementia (Prinz et al., 1982), and have recently been reported already in the stage of mild cognitive impairment (MCI; Hita-Yañez et al., 2013; Westerberg et al., 2012), often a precursor of dementia due to AD (Sperling et al., 2011). While decline in sleep quality is also common in healthy aging, including sleep parameters relevant for memory consolidation (Backhaus et al., 2007; Mander et al., 2013, 2014), the severity of the decline is significantly accelerated in patients with MCI or dementia due to AD (Prinz et al., 1982; Westerberg et al., 2012). Sleep disruptions not only contribute to memory deteriorations in MCI patients (Westerberg et al., 2012), but may also play a direct role in the progression of the underlying pathology (Wang et al., 2011; Ju et al., 2014).

Sleep plays an active role in long-term consolidation of memories (Diekelmann and Born, 2010). Specifically, slow oscillations (SO, large amplitude waves <1Hz) and sleep spindles (8-15 Hz), that can be measured by electroencephalography (EEG), were shown to be of particular importance for declarative memories (Schabus et al., 2004; Marshall et al., 2006; Wilhelm et al., 2011; Ngo et al., 2013). According to the “active system consolidation” account, newly encoded memories are reactivated during sleep, accompanied by sharp-wave ripple events (80-100 Hz) in the hippocampus, and redistributed to cortical long-term storage networks through a coordinated dialog between the hippocampus and neocortex (Rasch and Born, 2013). This dialog is mediated by a particular coupling between cortical SO and thalamo-cortical fast spindles (12-15 Hz) – with spindles preferably occurring during SO up-phases (Mölle et al., 2011; Ngo et al., 2013; Cox et al., 2014; Staresina et al., 2015) – and hippocampal ripples grouped at the troughs of fast spindles (Clemens et al., 2007; Staresina et al., 2015). Slow spindles (8-12 Hz) are a separate kind of sleep spindle activity whose function in memory consolidation is less well understood.

Apart from the consolidation aspect, there is mounting evidence that sleep, in particular SO, further promotes the clearance of cortical amyloid-β (Xie et al., 2013), a peptide involved in the pathogenesis of AD (Hardy and Selkoe, 2002). Strongest associations between SO activity and amyloid-β levels were found in the medial prefrontal cortex (Mander et al., 2015), a brain area in which accumulation of amyloid-β is evident particularly early in the pathogenesis of AD (Sepulcre et al., 2013).

Thus, interventions targeting sleep parameters may provide a therapeutic approach not only for memory consolidation deficits, but also to tackle the progression of Alzheimer pathology in MCI patients (Ju et al., 2014; Mander et al., 2016), with SO activity as promising target candidate. Application of slow oscillating weak transcranial direct current stimulation (so-tDCS, with frequency <1 Hz) during sleep provides a non-invasive method to enhance SO activity, as demonstrated in healthy young (Marshall et al., 2006) and older individuals (Westerberg et al., 2015; Ladenbauer et al., 2016; Paßmann et al., 2016) as well as in patients with attention deficit hyperactivity disorder (Prehn-Kristensen et al., 2014) or schizophrenia (Del Felice et al., 2015). In addition, this stimulation can lead to increased spindle activity (Marshall et al., 2006; Ladenbauer et al., 2016; Paßmann et al., 2016), and, importantly, to improved retention of declarative memories after night-time sleep (Del Felice et al., 2015; Marshall et al., 2006; Prehn-Kristensen et al., 2014), or a day-time nap (Westerberg et al., 2015; Ladenbauer et al., 2016).

Whether these beneficial effects of so-tDCS carry over to patients with MCI is unclear, and nontrivial, given severe impairments of sleep and memory as well as disruption of sleep-promoting brain structures in MCI (Ju et al., 2014). Furthermore, it is not known how this stimulation alters the cross-frequency coupling between SO and fast spindles, which is considered pivotal in memory consolidation during sleep. Here we addressed these questions by applying so-tDCS during an afternoon nap in patients with MCI and assessing its impact on relevant sleep characteristics, including cross-frequency coupling, as well as memory consolidation. Thereby we aimed to advance the potential of this noninvasive intervention towards therapeutic purposes, and provide novel insight on the underlying mechanisms.

## Results

We analyzed EEG data and memory performance of 16 MCI patients (7 female, mean age 70.6 years ± 8.9 SD) who were tested on memory tasks before and after a 90-min nap with either so-tDCS or sham stimulation (two sessions, balanced cross-over design). A schematic diagram of the experiment is shown in Figure 1. We focused on visual recognition memory, whose impairment occurs early in the course of AD (Barbeau et al., 2004) and thus constitutes a sensitive target for interventional approaches in MCI. For comparison with previous studies we additionally assessed word pair recall.

**Figure 1.**
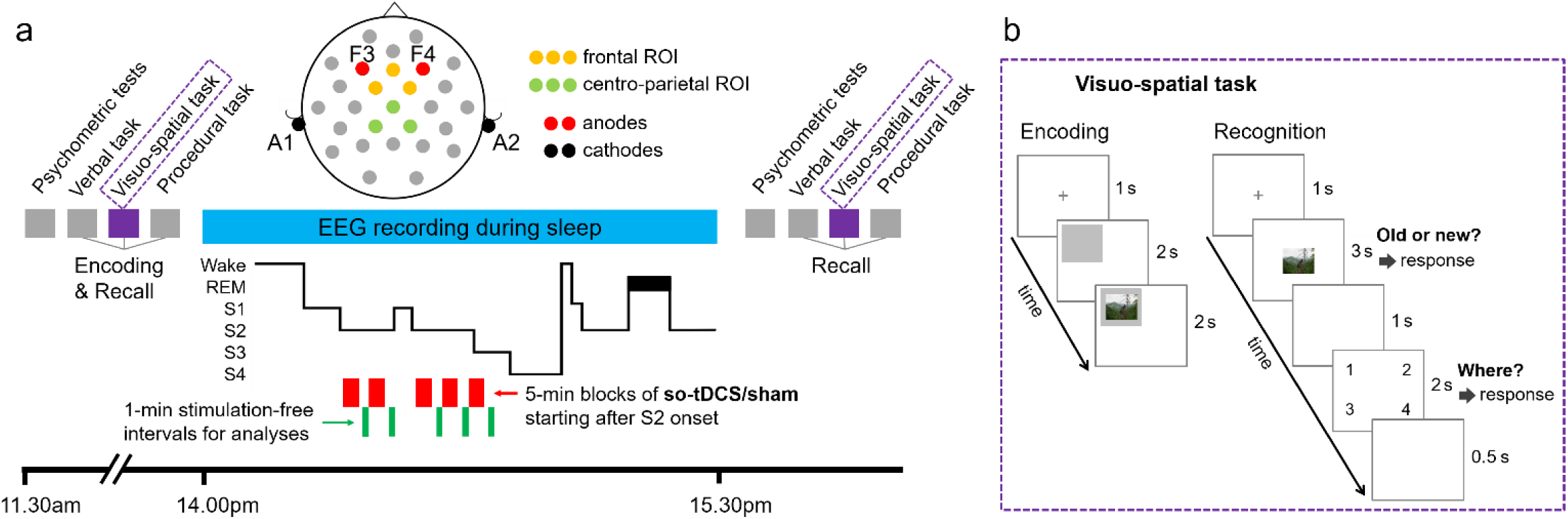
Study design. a) Subjects learned a verbal, a visuo-spatial and a procedural task in the indicated order following psychometric control tests. During a subsequent 90-min nap (14:00h to 15:30h) EEG was recorded and either so-tDCS or sham stimulation was applied (within-subject design, randomized order) in up to five 5-min blocks (red bars) which started after sleep stage 2 onset. According to the example hypnogram stimulation blocks are discontinued as the subject moves from sleep stage 2 to stage 1, and resumed after the subject again enters sleep stage 2 (or lower). REM, rapid eye movement sleep (black bar in the hypnogram); S1-S4, sleep stages 1-4. Recordings from electrodes Fz, FC1, FC2 (frontal region of interest, ROI) and Cz, CP1, CP2 (centro-parietal ROI) during 1-min stimulation-free intervals (green bars) starting 40 s after each so-tDCS/sham block were used for spectral and phase amplitude coupling analyses. Memory retrieval and psychometric control tests were performed 30 min after the nap. b) Example encoding and recognition trials of the visuo-spatial memory task. For encoding a gray rectangle was presented at one of the quadrants of the screen following a fixation cross and followed by a neutral picture within the gray region for 2 s. For recognition a picture was displayed in the center of the screen for 3 s following a fixation cross. Within this time period subjects were asked to indicate whether they believed had seen the picture earlier. If subjects recognized an item, they also indicated in which quadrant they believed the item had been presented. so-tDCS = slow oscillatory transcranial direct current stimulation.

We first asked whether so-tDCS beneficially affects memory-relevant sleep measures. For this purpose we focused on the immediate effects on SO and fast spindles measured by EEG during 1-min lasting stimulation-free intervals of up to five stimulation/sham blocks. We assessed the power of these oscillations as well as their functional coupling. Based on previous studies we performed these analyses for frontal and centro-parietal regions of interest separately (Mölle et al., 2011; Ladenbauer et al., 2016). In addition, we assessed frontal slow spindle power and quantified changes of sleep architecture. We then examined how stimulation changed declarative memory performance and finally explored the relationship between changes of performance and sleep measures.

The number of so-tDCS/sham blocks could vary between sessions, as we accounted for the individual subject’s sleep by ensuring sleep stage 2 or deeper prior to every stimulation block (state-dependent stimulation, see Methods). No significant differences were seen between conditions regarding the number of so-tDCS/sham blocks participants received (see Table 1, Wilcoxon signed-rank test: p = 0.453). The stimulation was well tolerated by all patients. They did not report any sensation except one patient who indicated a tingling perception. Post experimental debriefing indicated that most subjects were not able to guess in which nap session so-tDCS was applied (n=14 answered “do not know” while 2 subjects correctly guessed the so-tDCS nap). Considering sleep schedules and sleep duration prior to the experiments no differences between stimulation conditions were evident (all p’s > 0.2).

**Table 1.** so-tDCS/sham interval count and sleep architecture

|  | so-tDCS |  | sham |  |  |
| --- | --- | --- | --- | --- | --- |
| Distribution of stimulation counts during the nap |  |  |  |  |  |
| 5 blocks [n] | 12 |  | 10 |  |  |
| 4 blocks [n] | 2 |  | 3 |  |  |
| 3 blocks [n] | 2 |  | 3 |  |  |
|  | mean (SD) | MD | mean (SD) | MD | p |
| Sleep time excluding movement periods and so-tDCS/sham [min] | 48.8 (13.2) | 49.5 | 51.6 (11.6) | 56.00 | .569<sup>#</sup> |
| Proportion of total sleep time (including so-tDCS/sham) spent in different sleep stages |  |  |  |  |  |
| WASO [%] | 14.7 (11.9) | 11.3 | 20.36 (1) | 15.1 | .218 |
| NREM stage 1 [%] | 15.1 (5.7) | 11.8 | 16.6 (5.1) | 14.1 | .959 |
| NREM stage 2 [%] | 27.9 (6.7) | 28.9 | 24.4 (7.8) | 22.8 | .258 |
| NREM stage 3 [%] | 2.7 (1.1) | 0.3 | 2.7 (1.7) | 0.0 | .504<sup>#</sup> |
| NREM stage 4 [%] | 1.1 (1.1) | 0.0 | 0.2 (0.1) | 0.0 | 1.000<sup>#</sup> |
| REM [%] | 2.3 (1.3) | 0.0 | 3.1 (1.3) | 0.0 | .600<sup>#</sup> |
| Proportion of time during stimulation-free intervals spent in different sleep stages |  |  |  |  |  |
|  | mean (SD) | MD | mean (SD) | MD | p |
| WASO [%] | 7.44 (10.5) | 0.0 | 14.7 (15.0) | 11.7 | .062<sup>#</sup> |
| NREM stage 1 [%] | 7.5 (11.6) | 3.3 | 15.8 (19.9) | 11.7 | .091<sup>#</sup> |
| NREM stage 2 [%] | 70.3 (21.0) | 69.5 | 51.5 (18.4) | 53.3 | .021 |
| NREM stage 3 [%] | 12.8 (11.9) | 10.0 | 10.4 (14.2) | 4.9 | .451<sup>#</sup> |
| NREM stage 4 [%] | 2.8 (9.5) | 0.0 | 3.1 (9.5) | 0.0 | 1.000<sup>#</sup> |
| REM [%] | 0.0 (0.0) | 0.0 | 2.3 (6.3) | 0.0 | .180<sup>#</sup> |

Sleep time = elapsed time from sleep onset to the end of nap; WASO = wake after sleep onset; SD = standard deviation; MD = median. Comparisons are based on paired-samples t-test unless indicated otherwise. \* $p < 0.05$ . <sup>#</sup> Wilcoxon signed-rank test due to skewed distributions.

### Effects on memory-relevant sleep measures

#### Spectral power

To analyze the effects of stimulation on spectral power memory-relevant frequency bands we used a linear mixed model (LMM) which accounts for individual differences in sleep physiology and unequal numbers of observations across subjects (3-5 so-tDCS/sham blocks, see Methods). LMM analyses revealed a significant so-tDCS related enhancement in frontal as well as centro-parietal SO (0.5-1 Hz) power during the nap (*stimulation,* mean difference ± standard error, frontal: 0.279 ± 0.06, p < 0.001; centro-parietal: 0.185 ± 0.066, p = 0.006), see Figure 2. This enhancement could not be explained by the baseline SO power (during 1 min. prior to the first so-tDCS/sham block) for which no significant difference between so-tDCS and sham condition was observed (frontal: t_(15)_ = 1.65, p = 0.121; centro-parietal: t_(15)_ = 1.20 p = 0.249). Furthermore, baseline SO power was not significantly associated with SO activity following the stimulation blocks (*baseline,* frontal: *β_2_* = −0.082, standard error (SE) = 0.168, p = 0.628; centro-parietal: *β_2_* = −0.145, SE = 0.206, p = 0.483). With regard to the course of SO power over 1-min stimulation-free intervals across the nap independent of condition, we did not observe a significant linear change *(time,* frontal: *β_3_* = 0.027, SE = 0.029, p = 0.353; centro-parietal: *β_3_* = 0.027, SE = 0.033, p = 0.408), but an inverted U-shaped relationship, which was stronger over centro-parietal *(time^2^, β_4_* = −0.061, SE = 0.027, p = 0.025) than frontal sites *(time^2^, β_4_* = −0.040, SE = 0.024, p = 0.092). Neither the slope nor the curvedness of SO power over intervals significantly differed between conditions (*time×stimulation* and *time^2^×stimulation* interactions: all p’s > 0.2).

This enhancing so-tDCS effect was not restricted to the SO band. For fast spindle power (12-15 Hz), LMM analyses also showed significant stimulation effects (*stimulation,* mean difference ± standard error, frontal: 0.139 ± 0.034, p < 0.001; centro-parietal: 0.072 ± 0.035, p = 0.41) indicating increased power following so-tDCS as compared to sham. These so-tDCS-induced power increases could also not be explained by fast spindle power during baseline as there was no significant difference between so-tDCS and sham during baseline (frontal: t_(15)_ = 1.84, p = 0.086; centro-parietal: t_(15)_ = 0.88 p = 0.393). Nevertheless, we found that baseline fast spindle power was significantly associated with power in this frequency range during later stimulation free intervals over frontal (*baseline, β_2_* = 0.463, SE = 0.122, p < 0.001) and centro-parietal sites *(baseline, β_2_* = 0.392, SE = 0.146, p = 0.010).

**Figure 2.**
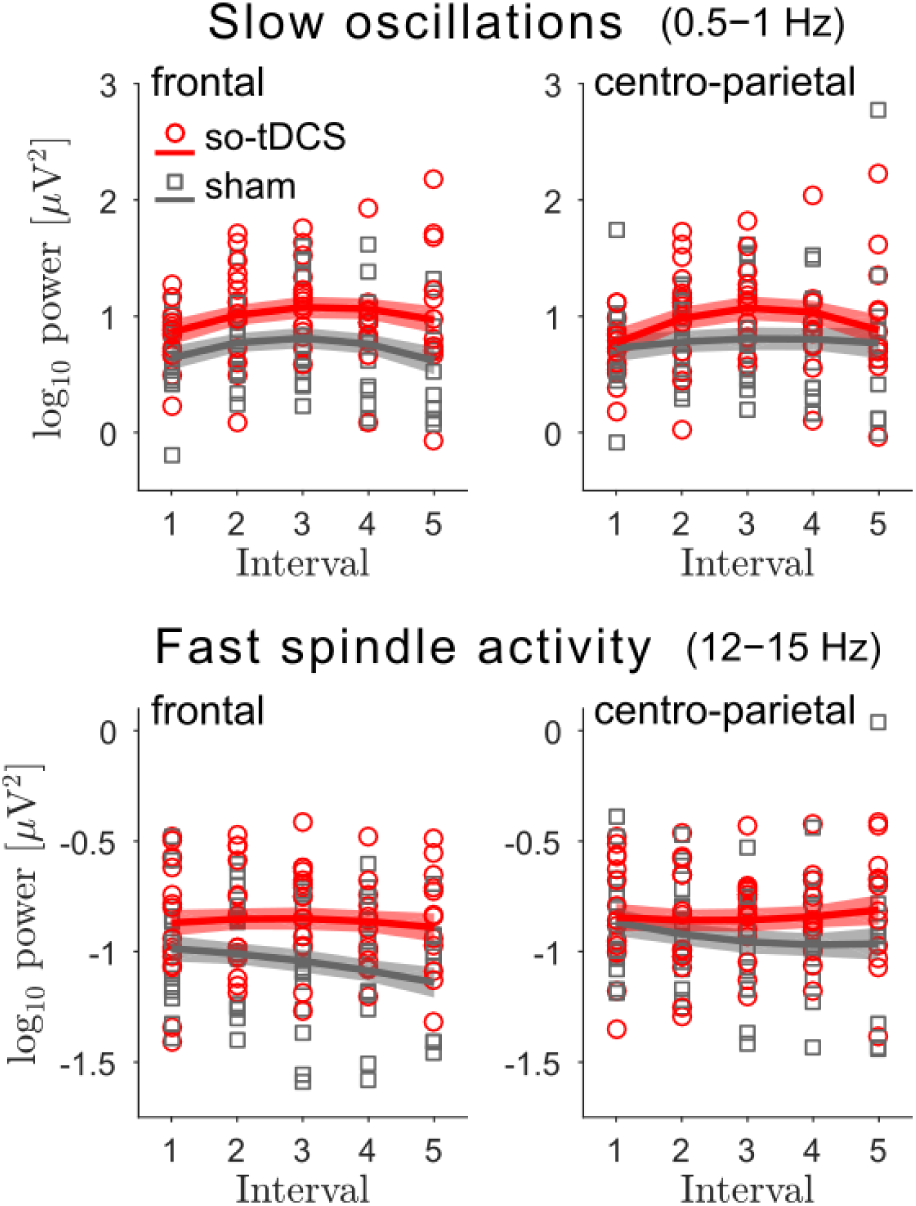
so-tDCS enhances EEG power in the slow oscillation and fast spindle frequency ranges. EEG power in the slow oscillations (0.5-1 Hz, top) and fast spindle (12-15 Hz, bottom) frequency ranges for the five 1-min stimulation-free intervals, for so-tDCS (red) and sham (gray) condition and considering the frontal (left) and centro-parietal (right) region of interest. Mean estimates ± SEM (shaded regions) from the linear mixed model (LMM) are included.

We further examined the effect of stimulation on frontal slow spindle power (8-12 Hz), as a previous study in young subjects found so-tDCS-induced increases in the slow spindle band but not for fast spindle power (Marshall et al., 2006). In the present study, frontal slow spindle power was significantly increased following so-tDCS compared to sham stimulation (*stimulation,* mean difference ± standard error, 0.101 ± 0.031, p = 0.001). Here, baseline frontal spindle power did not significantly differ between so-tDCS and sham condition (t_(15)_ = 1.11, p = 0.287), but in line with fast spindle power, baseline slow spindle power was significantly associated with power in this frequency range during stimulation free intervals following so-tDCS/sham blocks *(baseline, β_2_* = 0.674, SE = 0.112, p < 0.001). No further significant effects for power in the fast and slow spindle frequency ranges were found (all p’s > 0.1).

#### Phase-amplitude coupling

To assess functional coupling of SOs and spindles, we employed an analytical approach which asked whether the amplitude (power) of the spindle oscillation is systematically modulated by the phase of the SO (phase-amplitude coupling, PAC). Using an event-locked analysis we specifically examined the hypothesized nesting of SOs and fast spindles. We identified SO events in the EEG according to an established detection algorithm and aligned time-frequency representations (TFRs) of the peri-event epochs to the center of the SO trough (see Methods). Thereby SO-to-spindle PAC emerges as power modulation over time in the respective (event-locked) TFR. To ensure reliable PAC results four patients had to be excluded from this analysis because they exhibited only a very small number of SO events (<10 per electrode). Thus, n=12 were included in the final analysis.

We found that so-tDCS lead to a significant increase in centro-parietal fast spindle power during the SO up-phase that follows the event-centering SO trough (down-phase) as well as in centro-parietal slow spindle power during the SO up-phase that precedes the SO trough, see Figure 3a-c. It may be noted that the modulation of slow spindle power in the so-tDCS condition led to visibly stronger oscillatory behavior which was phase delayed by about 250 ms relative to the fast spindle power oscillation (Figure 3c). In the frontal region of interest we did not observe significant effects of stimulation on PAC assessed in this way.

**Figure 3.**
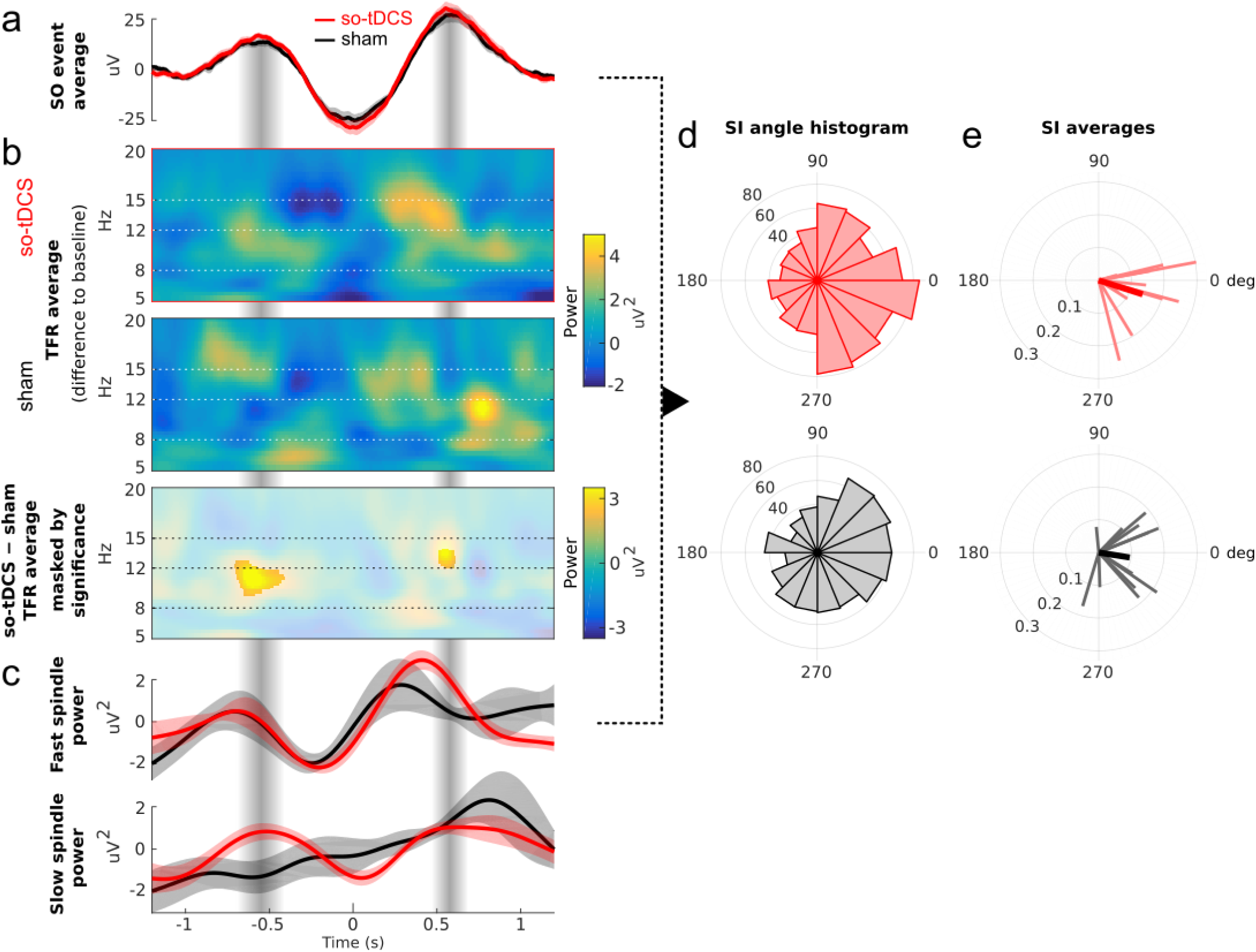
Phase amplitude coupling between SO and spindle power. a) Grand average EEG trace (mean ± SEM across participants) of a total of 860 events for so-tDCS (red) and 765 events for sham (black) condition aligned to the slow oscillations (SO) trough (time 0), from the centro-parietal ROI and all 1-min stimulation-free intervals. b) Time-frequency representations (TFRs) locked to the SO events (from a) and averaged per condition: so-tDCS (top) and sham (center). Shown are the differences to the pre-event baseline power values (−2.5 s to −1.2 s). Below: difference of these TFRs, masked by significance (bottom, p < 0.05, corrected). c) Time course of event-locked average power from the TFRs in b), filtered in the range of the modulating SO, for the fast spindle (12-15 Hz, top) and slow spindle (8-12 Hz, bottom) frequency ranges (mean ± SEM across participants). d) Histogram of synchronization index (SI) angles indicating the phase difference between SO and the fast spindle power fluctuation [cf. a) and c)] for the conditions so-tDCS (top, n=860) and sham (bottom, n=765). An angle value of 0 indicates synchrony, whereas 180 deg. indicates an anti-phase relationship. A value just below 0 (close to 360 deg.) indicates that spindle power tended to peak shortly before the SO peak. Note that the SI angle distribution for so-tDCS indicates that the spindle power peak preferably occurred during the late rising phase of SO. e) SIs averaged per subject (thin lines) and across subjects (thick lines) for the two conditions. Note that the angle of the SI indicates the phase of SO at which spindles tend to occur, whereas its radius indicates the strength of locking (coupling) between SO and the oscillatory spindle power fluctuations.

**Figure 4.**
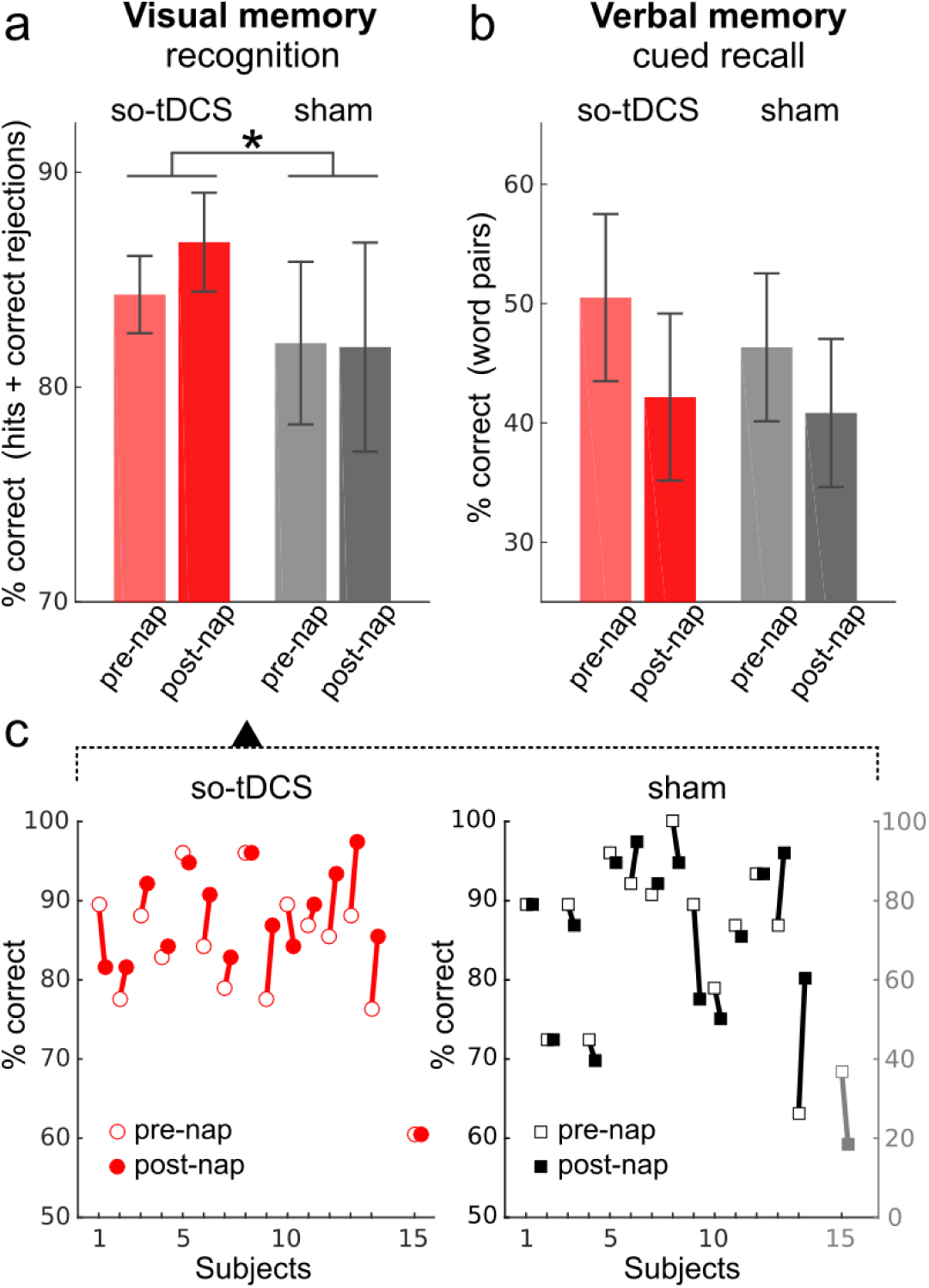
Retention performance in declarative memory tasks in the so-tDCS versus sham condition. a) Recognition performance (percent correct: proportion of hits and correct rejections) in the picture memory subtask and b) cued recall performance (percent correct) in the verbal memory task for so-tDCS (red) and sham stimulation (gray), measured before (pre-nap) and after (post-nap) the nap. A significant stimulation effect emerged for picture memory, with higher picture recognition performance following so-tDCS compared to sham condition. Data are expressed as means ± SEM; *p < 0.05. c) Picture recognition performance of individuals before and after napping for so-tDCS (left) and sham (right) condition. Note the separate scale for the outlier (subject 15, gray) in sham.

We further examined this coupling between SO and fast spindles in an event-wise manner using a synchronization index (SI) which measures the strength of locking and phase shift between the modulating SO and the modulated oscillatory spindle power fluctuation (see Methods). The SIs shown in Figure 3d and 3e emphasizes the enhancement of the hypothesized memory-relevant PAC due to stimulation. The overall distribution of SI angles indicates the phases of SO at which fast spindles occur preferably. This distribution shows a much stronger peak close to zero (in phase synchrony) for stimulation compared to sham condition (Figure 3d) and is more concentrated as measured by the resultant vector length (so-tDCS: 0.20 vs. sham: 0.16) and a larger proportion of SI angles are in the fourth quadrant (270-360 deg., so-tDCS: 35% vs. sham: 29%), indicating that spindle power peak preferably occurred during the late rising phase of SO. This was confirmed by the statistics for the subject-averaged SI angles (Figure 3e), for which 8 (so-tDCS) vs. 5 (sham) out of 12 are in the fourth quadrant and whose resultant vector length (indicating the degree of concentration across subjects) amounts to 0.89 (so-tDCS) vs. 0.58 (sham). V-test results further demonstrate that for the stimulation condition subject-averaged SI angles are substantially tighter concentrated close to 0 (so-tDCS: V=10.16, p<.00002 vs. sham: V=6.84, p<.003).

For the frontal region of interest we observed a similar (but weaker) enhancement of SO-to-fast-spindle PAC by stimulation as quantified by V-test statistics for the subject-averaged SI angles (so-tDCS: V=8.50, p<.0003 vs. sham: V=5.67, p=.01) and corresponding resultant vector lengths (so-tDCS: 0.71 vs. sham: 0.52).

#### Sleep stages

Table 1 summarizes total sleep time spent in different sleep stages during the nap and during the 1-min stimulation-free intervals. No significant differences between conditions were found in total sleep time and times spent in the different sleep stages (all p’s > 0.2). However, for the 1-min stimulation-free intervals pairwise comparisons yielded a significant difference in non-rapid eye movement (NREM) sleep stage 2 (t_(15)_ = 2.57, p = 0.021). Sleep stage 2 (in %) was increased following so-tDCS as compared to sham, while wake time after sleep onset and NREM sleep stage 1 was trendwise reduced (Wilcoxon signed-rank tests: p = 0.062 and p = 0.091, respectively).

To summarize, so-tDCS in MCI patients profoundly increased SO power and enhanced power in the fast and slow spindle frequency ranges. PAC analyses further revealed that so-tDCS lead to stronger synchronization between SO and fast spindle power, in particular by increasing spindle power during late rising SO up-phases. These beneficial so-tDCS effects were also reflected in the sleep structure during the 1-min stimulation-free intervals, with increased sleep stage 2 and trendwise less sleep stage 1 and wake phases as compared to sham condition.

### Effects on memory

To control for potential confounding influences on memory performance we assessed selfreported mood, activation, sleepiness and attention before and after the naps. No significant pre- and post-nap differences were found (all p’s > 0.1). Likewise we did not find stimulation-dependent changes in attention, mood and activation (all p’s > 0.2), but a nonsignificant trend toward increased self-reported sleepiness as indexed by the Tiredness Symptoms Scale (physical sleepiness) following naps with so-tDCS (F_(1,15)_ = 3.50, p = 0.082) compared to sham condition. We tested whether so-tDCS effects were also reflected in visual memory performance in MCI patients (n=15, see Methods). Given the (mild) so-tDCS effect on sleepiness, we included sleepiness as covariate in the respective analyses on memory performance (analysis of covariance for repeated measures, ANCOVA). We found a significant so-tDCS effect on visual memory as assessed by picture recognition accuracy (percent correct sores: hit rate + correct rejection rate) (STIMULATION X TIMEPOINT interaction: F_(1,12)_ = 5.34, p = 0.039, performance change across nap in %: so-tDCS, 2.89 ± 1.35; sham, −0.96 ± 2.11). We further tested for pre-nap differences between conditions and possible response bias. Pre-nap visual recognition performance did not differ between sessions (t_(14)_ = 0.38, p = 0.712; see also Figure 5). With regard to response bias, as indicated by the sum between hit rate and false alarm rate, an impact towards less conservative responding after the nap with so-tDCS as compared to sham condition was found (F _(1,12)_ = 14.55, p = 0.002). However, examining this effect in more detail by testing so-tDCS effects separately for hit rate and false alarm rate, a significant effect was only apparent for hit rate (F_(1,12)_ = 5.91, p = 0.002), but not for false alarm rate (F_(1,12)_ = 2.54, p = 0.137). Additionally, we evaluated the so-tDCS effect on visual memory without correcting for sleepiness in the analysis (rmANOVA, repeated measures analysis of variance). Here, stimulation-induced effect failed to reach significance, but a statistical trend in favor of stimulation was noted (F_(1,14)_ = 3.58, p = 0.079). With regard to main effects of STIMULATION and TIMEPOINT on visual recognition performance, no significant effects were evident, but a trend for a main effect of session emerged (STIMULATION, F_(1,12)_> = 3.80, p = 0.075).

**Figure 5.**
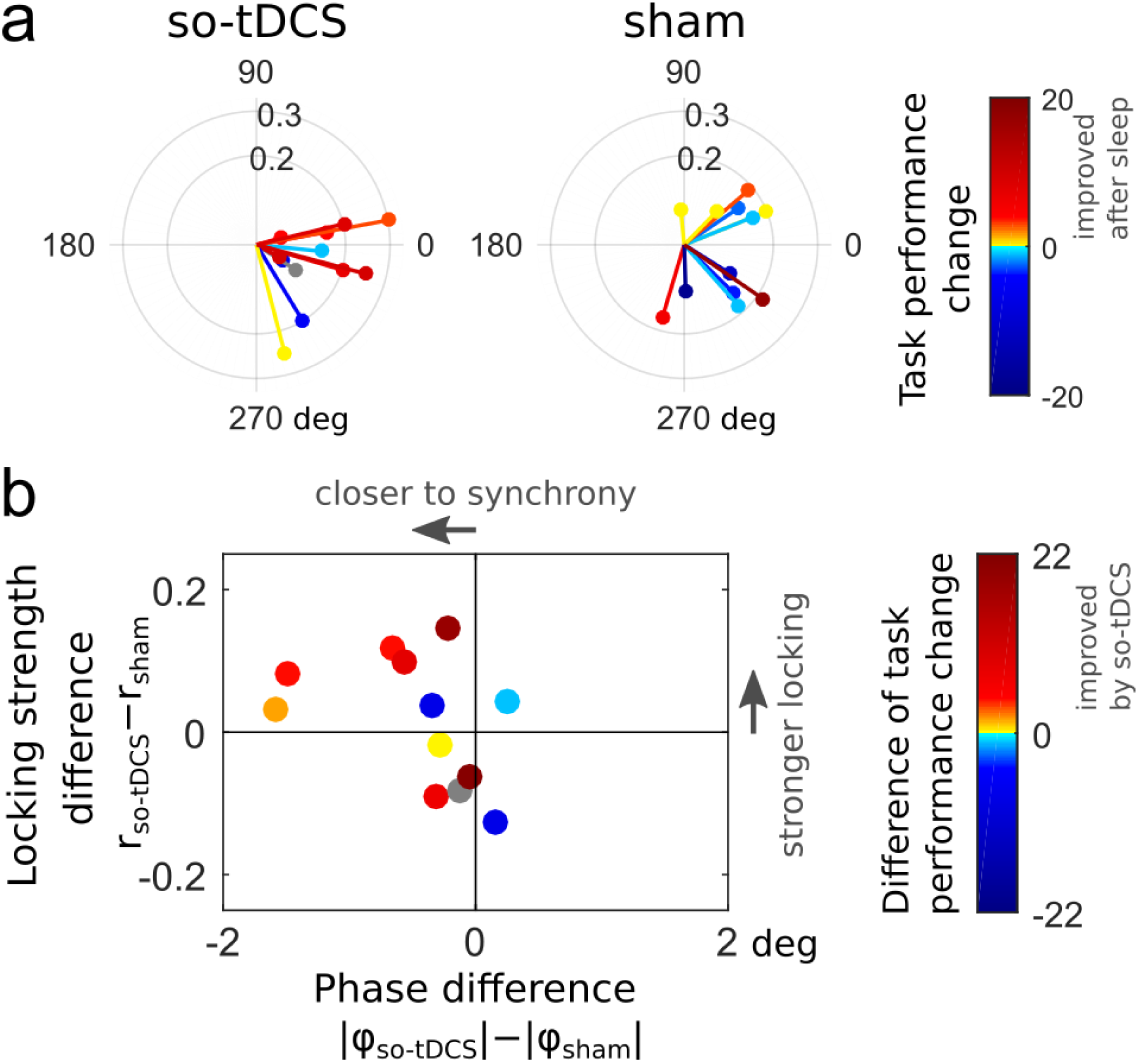
Relationship between SO-to-fast spindle phase amplitude coupling and visual memory task performance. a) Subject-averaged synchronization index (SI) values from the centro-parietal region of interest (cf. Figure 4b) colored according to the change in visual task performance (post-nap - pre-nap) for so-tDCS (left) and sham (right) condition. Warm color indicates an improvement after the nap. b) Subject-wise difference of SI radii between conditions (indicating the change in locking strength) versus difference of absolute SI angles (reflecting the change in “preferred” SO phase at which fast spindles tend to occur), colored according to the difference of visual task performance change (“so-tDCS - sham”). Warm color indicates an improvement due to so-tDCS.

The accuracy in location memory did not show any so-tDCS-related performance change (STIMULATION x TIMEPOINT interaction: F_(1,12)_ = 0.01, p = 0.913; performance change across nap in %: so-tDCS, −8.37 ± 4.21 sham, −4.95 ± 5.92), but a decline of spatial memory following the nap independent of stimulation condition (main effect TIMEPOINT: F_(1,12)_ = 6.18, p = 0.029; pre-nap in %: −12.87 ± 11.72, post-nap in %: −19.53 ± 10.19).

For comparison with previous studies, we also examined so-tDCS effects on retention performance in the word paired-associate learning task (verbal memory). We observed no so-tDCS effect on cued recall performance (statistically controlling for sleepiness, STIMULATION x TIMEPOINT interaction: F_(1,12)_ = 2.27, p = 0.156, n=15), but MCI patients showed a significant decrement of recall performance independent of stimulation condition from pre- to post nap (main effect TIMEPOINT: F_(1,12)_ = 12.96, p = 0.003; pre-nap in %: 45.9 ± 6.6; post-nap in %: 39.1 ± 6.1). Baseline verbal memory performance before napping did not differ between so-tDCS and sham session (t_(14)_ = 1.03, p = 0.322).

To determine the specificity of so-tDCS effects, a procedural memory task was assessed in addition to declarative memory tasks. Neither main nor interaction effects were noted (all p’s > 0.5).

In sum, so-tDCS effects on the behavioral level revealed beneficial effects on the recognition performance of pictures after correcting for the confounding variable sleepiness. No effects of so-tDCS were seen on the location memory subtask, the verbal memory and procedural memory task.

### Relationship between effects on sleep measures and memory performance

To examine the relationship between stimulation-induced changes in sleep measures (SO and spindle power changes as well as sleep stages) and visual memory performance changes, we correlated differences in sleep measures between stimulation conditions with the difference in visual recognition (pre- to post-nap) performance change between so-tDCS and sham condition. For power measures as well as sleep stages no significant correlations were found (all p’s > 0.3). Finally, we asked whether coupling between centro-parietal SO and fast spindles, measured by the SI, was related to visual task performance change. We found that a positive “post-nap - pre-nap” task performance difference tended to be associated with synchronized SO-to-fast-spindle PAC in so-tDCS condition (SI angles close to 0), while no such relationship could be observed for sham condition (Figure 5a).

Considering the relationship between stimulation-induced changes in PAC and visual recognition performance, an improvement in memory performance due to so-tDCS tended to accompany stronger synchronization (SI phase) between SO and fast spindle power, when taking both phase differences (SI angles) and locking strengths (SI radii) into account (Figure 5b). Fast spindle power during the SO up-phase (after the event-centering down-phase) has previously been found to correlate with (verbal) memory performance (Ngo et al. 2013); therefore we additionally applied a very similar measure (fast spindle power peak and its timing during that SO up-phase) but found no correlation with visual task performance change.

To summarize this section, stimulation-induced improvement of visual memory did not significantly correlate with changes in overall SO or spindle power, but was rather associated with enhanced synchronization between SO and fast spindle power fluctuations.

## Discussion

In light of the mounting evidence for the association between sleep disruptions and memory decline in MCI, we investigated whether specific memory-relevant sleep characteristics - in particular, SO, fast spindles and the coupling between these oscillations - could be enhanced by so-tDCS during a daytime nap in MCI patients. Moreover, we intended to promote visual memory consolidation via stimulation in this patient group. We showed, for the first time in MCI patients, that so-tDCS significantly increased SO and spindle power. Importantly, we demonstrated that so-tDCS led to an enhanced endogenous SO-to-spindle coupling in the following way: spindle power was significantly amplified during the depolarizing SO up-phases and synchronization between SO and the fast spindle power signal was stronger. In addition we found an improvement in visual memory performance which tended to accompany stronger synchronization between SO and fast spindle power.

### so-tDCS increases SO and spindle activity as well as their functional coupling in MCI patients

The impact of so-tDCS on SO power in the present study is well in line with previous studies examining the effects of so-tDCS during sleep on memory-relevant sleep measures in healthy older (during nighttime sleep: Paßmann et al., 2016; during daytime nap: Ladenbauer et al., 2016; Westerberg et al., 2015) and healthy young adults (during nighttime sleep, Marshall et al., 2006). Our observation of amplified fast spindle activity over frontal and centro-parietal sites is consistent with previous results from our group in older adults (Ladenbauer et al., 2016; Paßmann et al., 2016) but contrary to those of Westerberg et al. (2015) in healthy older and of Marshall et al. (2006) in healthy young individuals. A possible reason for this discrepancy may be the variation in stimulation procedure, as studies from our group controlled for ongoing sleep (sleep stages 2,3 or 4) prior to every stimulation block (state-dependent stimulation, in contrast to Westerberg et al. 2015). Since stimulation effects strongly depend on ongoing brain state (Marshall et al., 2011) a sleep state-dependent protocol of this kind may be critical, specifically for older adults, given their increased sleep fragmentation (Bliwise et al., 2009), an issue potentiated in patients with neurodegenerative disease (Lim et al., 2013).

Considering the coupling between SO and spindles we observed in both conditions that SO modulated fast spindle power such that it exhibited up- and down-phases similar to SO on average, with positive peaks occurring during the depolarizing SO up-phase close to (but slightly before) the SO peak (cf. Figure 3c). These stimulation-independent observations are well in line with previous results (Mölle et al., 2002, 2011; Niknazar et al., 2015; Staresina et al., 2015). Furthermore, synchronization between SO and fast spindle power in sham condition as quantified by subject-averaged synchronization indices is comparable to what was reported in Staresina et al. (2015) (V=6.84 here vs. V=6.55 there), who demonstrated coupling between SO and spindles as well as between spindle and ripple activity in a hierarchical manner in humans. Notably, we provided evidence for a strong enhancement of SO-to-spindle coupling by so-tDCS – quantified by both fast spindle power during SO up-phases and synchronization (cf. Figure 3b-d). This is a particularly promising result in the light of accumulating evidence for the functional role of that specific coupling (Ruch et al., 2012; Ngo et al., 2013; Niknazar et al., 2015).

### so-tDCS improves visual but not verbal declarative memory in MCI patients

We found that so-tDCS led to improved visual recognition performance, consistent with our previous study on healthy older individuals (Ladenbauer et al., 2016). MCI patients showed an even larger performance gain due to so-tDCS on average (MCI: mean gain=3.86, SD=7.90; healthy older: mean gain=2.48, SD=3.66) but also the variances were larger. Regarding location memory (retrieval of picture locations) and verbal memory (word pair task) we found no effect of so-tDCS in MCI patients, similar to the results in healthy older adults (Ladenbauer et al., 2016).

For the spatial subtask this may be due to the small number of valid items (retrieval of location only if participant recognized the picture as “known”) or the difficulty of the task for our participants in the light of recent evidence for beneficial so-tDCS effects primarily on consolidation of relatively simple information (Barham et al., 2016). For the verbal task relatively strong semantic associations in our word pairs (Drosopoulos et al., 2007; Ladenbauer et al., 2016) may have decreased the sensitivity of this task to detect stimulation-related memory effects.

### Relationship between changes of sleep characteristics and visual memory improvement

Changes in overall SO or spindle power were not significantly correlated with visual performance change, similar to previous so-tDCS studies that could not show a correlation between SO enhancement and memory improvement (Marshall et al., 2006; Prehn-Kristensen et al., 2014; Del Felice et al., 2015; Westerberg et al., 2015; Ladenbauer et al., 2016). This may be attributable to low statistical power because of small sample sizes. Each of these oscillations on their own (without intervention) have been correlated with overnight retention of memories (Schabus et al., 2004; Tamminen et al., 2011; Mander et al., 2014), however, a recent study with large sample size (n=929) challenges these results (Ackermann et al., 2015). It is well likely that the functional coupling between SO and spindles plays a more dominant role for memory consolidation. From our results stimulation-induced improvement in visual memory is predictable by enhanced synchronization between SO and fast spindle power (Figure 5b). Nevertheless, it should be noted that in-phase synchrony (SI angles = 0) may not necessarily be the optimal relationship for memory consolidation as compared to a (small) phase difference instead (e.g., fast spindle power peak shortly before SO peak, see Cox et al., 2014; Staresina et al., 2015). Future studies should determine whether stronger SO-to-spindle coupling (in synchrony or locked with a particular phase shift) consistently leads to improved consolidation during sleep.

Our results are very encouraging in the light of strong evidence for sleep deteriorations and their consequences on cognitive functions in MCI: sleep disturbances are associated with a decline in memory consolidation (Westerberg et al., 2012), facilitate accumulation of amyloid-β (Kang et al., 2009), and may thereby trigger the pathophysiological process leading to AD (Ju et al., 2014; Mander et al., 2016). Thus, optimizing sleep by so-tDCS constitutes a promising therapeutic approach in two respects. First, enhancement of memory-relevant sleep parameters (SO, spindles and their functional coupling) can lead to improved memory consolidation, which helps to preserve cognitive abilities and possibly reduces clinical severity in AD (Ju et al., 2014). Second, improvement of sleep physiology, in particular SO activity, may decelerate the progression of disease pathology in MCI patients through enhanced clearance of amyloid-β (Xie et al., 2013). In these regards, so-tDCS may be more effective compared to pharmacological treatments, which so far could not enhance both SO activity and functional SO-to-spindle coupling during sleep at the same time (Feld et al., 2013; Niknazar et al., 2015), and which failed to improve, or even decreased memory performance (Vienne et al., 2012; Feld et al., 2013; Hall-Porter et al., 2014), with one exception (Niknazar et al., 2015).

### Limitations and Conclusion

A few limitations of the study should be noted: First, patients tended to feel sleepier following a nap with so-tDCS as compared to sham condition. This effect on sleepiness was also found in healthy older individuals in our previous study (which was significant), while no other study applying so-tDCS during sleep reported this effect before. We cannot rule out that increased SO activity due to so-tDCS persisted until the survey after the nap (Nitsche and Paulus, 2001),although it is very unlikely. Note, however, that we statistically controlled for sleepiness to exclude its potential effect on memory performance. Second, to analyze the immediate effects of so-tDCS on coupling between SO and spindles a rather limited amount of EEG data (from up to five 1-min stimulation-free intervals) could be used due to the study design. This explains why the power patterns in the TFRs – despite strong similarities – do not emerge as clear as in methodologically related studies, particularly in the slow spindle frequency range (e.g., Mölle et al., 2011; Staresina et al., 2015). However, the overall agreements of our results with previous ones support the validity of these analyses. Furthermore, we observed that visual recognition performance varied stronger across MCI patients in the sham as compared to so-tDSC session. Note however, that the study was balanced with respect to so-tDCS/sham order as well as task versions, and no other difference than stimulation condition occurred between sessions.

Despite these limitations, our results clearly demonstrate the potential of so-tDCS to enhance functional sleep physiology in MCI, including for the first time the coupling of SO to spindle activity – a mechanistic component considered crucial for the transfer of memories from hippocampus to cortical long-term storage networks. Apart from benefits on memory consolidation, as shown here for visual recognition memory, the stimulation may thereby delay the progression of Alzheimer’s pathology (Ju et al., 2014). Therefore it would be worthwhile to assess the extent to which this noninvasive stimulation method can be optimized for individualized treatment, for example, by fine-tuning to each patient’s sleep structure in an automated closed-loop manner (Ngo et al., 2013).

## Materials and Methods

This study was approved by the Ethics Committee of the Charité University Hospital Berlin, Germany, and was in accordance with the declaration of Helsinki. All subjects gave informed written consent before participation in the study and received a small reimbursement at the end.

### Participants

Twenty-two patients (10 female, mean age 71.2 years ± 8.79 SD; range: 50 - 81 years) with mild cognitive impairment (MCI) recruited from the memory clinic of the Department of Neurology of the Charité University Hospital Berlin participated in the present study. MCI patients (amnestic; single and multiple domain) were clinically diagnosed according to Mayo criteria based on subjective cognitive complaints and objective memory impairment in standardized tests. These reflected scores of at least 1 SD (thus, including both early and late MCI, see Jessen et al., 2014) below age- and education-specific norm in relevant subtest of the CERAD-Plus test battery (Memory Clinic, 2009; relevant subtests: Total Word List, Delayed Recall Word List/Figures) or the AVLT (German version of Auditory Verbal Learning Test; Helmstaedter et al., 2001) with relatively preserved general cognition, no impairment in activities of daily living, and no dementia (Sperling et al., 2011).

Exclusion criteria comprised MMSE<24 and history of severe untreated medical, neurological, and psychiatric diseases; sleep disorders; alcohol or substance abuse; brain pathologies identified in the MRI scan; intake of medication acting primarily on the central nervous system (e.g., antipsychotics, antidepressants, benzodiazepines, or any type of over-the-counter sleep-inducing drugs like valerian); and non-fluent German language abilities. Moreover, psychiatric comorbidity was monitored by Beck’s Depression Inventory-II (BDI-II, exclusion if BDI-scores ≥19; Kuehner et al., 2007) and Magnetic Resonance Imaging (MRI) of the brain was carried out to exclude major brain pathologies like brain tumor and previous stroke. Further, participants were excluded from the analysis if they failed to sleep long and deep enough for at least three so-tDCS/sham blocks, which corresponded to about 22 min spent in sleep stage 2 or slow wave sleep. This is due to our stimulation protocol, which was designed to prevent application of so-tDCS during inappropriate brain states (wake, sleep stage 1, or REM sleep).

In total, data from six patients had to be excluded due to insufficient sleep (n 2), inability to complete computerized tasks (n=3), and a previously undetected psychiatric disorder (n=1), resulting in a final group of 16 patients who completed the experiment. Excluded patients did not differ in baseline parameters from this cohort, apart from education duration (p = 0.048, all other p’s > 0.2; see Table 2 for Baseline characteristics), with higher mean education duration for excluded participants due to an outlier (30 years).

**Table 2.**
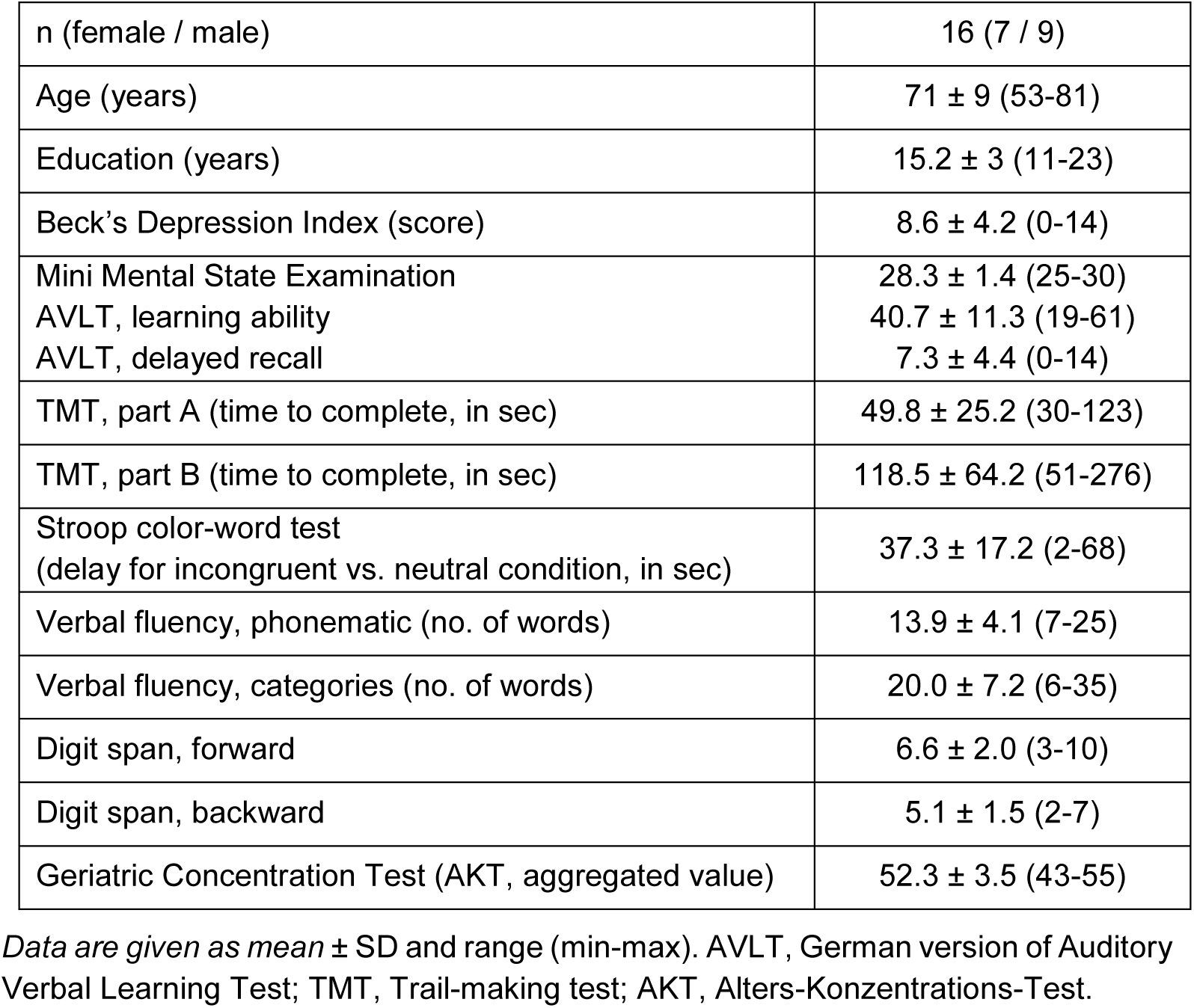
Baseline characteristics of subjects

|  |  |
| --- | --- |
| n (female / male) | 16 (7 / 9) |
| Age (years) | 71 ± 9 (53-81) |
| Education (years) | 15.2 ± 3 (11-23) |
| Beck's Depression Index (score) | 8.6 ± 4.2 (0-14) |
| Mini Mental State Examination | 28.3 ± 1.4 (25-30) |
| AVLT, learning ability | 40.7 ± 11.3 (19-61) |
| AVLT, delayed recall | 7.3 ± 4.4 (0-14) |
| TMT, part A (time to complete, in sec) | 49.8 ± 25.2 (30-123) |
| TMT, part B (time to complete, in sec) | 118.5 ± 64.2 (51-276) |
| Stroop color-word test<br>(delay for incongruent vs. neutral condition, in sec) | 37.3 ± 17.2 (2-68) |
| Verbal fluency, phonematic (no. of words) | 13.9 ± 4.1 (7-25) |
| Verbal fluency, categories (no. of words) | 20.0 ± 7.2 (6-35) |
| Digit span, forward | 6.6 ± 2.0 (3-10) |
| Digit span, backward | 5.1 ± 1.5 (2-7) |
| Geriatric Concentration Test (AKT, aggregated value) | 52.3 ± 3.5 (43-55) |

*Data are given as mean ± SD and range (min-max). AVLT, German version of Auditory Verbal Learning Test; TMT, Trail-making test; AKT, Alters-Konzentrations-Test.*

### Baseline assessments

A comprehensive neuropsychological testing for assessment of general cognitive status was administered to each participant comprising memory performance (German version of Auditory Verbal Learning Test (AVLT); Helmstaedter et al., 2001), working memory (digit span; Wechsler, 1997), executive functions (Stroop color-word test; Van der Elst et al., 2006), verbal fluency (Regensburg Verbal Fluency Test (RWT); Aschenbrenner et al., 2000), processing speed and set shifting (Trail Making test (TMT) part A and B; Tombaugh, 2004), selective attention and concentration (AKT; Gatterer, 2008). Furthermore, the neuropsychology battery developed by the Consortium to establish a Registry for Alzheimer’s Disease (CERAD-Plus; www.memoryclinic.ch) was administered. The affective state at the time of the testing was assessed using the Positive and Negative Affect Schedule (PANAS; Watson et al., 1988). For these baseline characteristics see Table 2.

In addition, questionnaires regarding recent sleep habits (the German version of Morningness-Eveningness-Questionnaire (d-MEQ); Griefahn et al., 2001), sleep quality (Pittsburgh Sleep Quality Index (PSQI); Buysse et al., 2015) and daytime sleepiness (Epworth Sleepiness Scale (ESS); Johns, 1991), and the Essen questionnaire on age and sleepiness „Essener Fragebogen Alter- und Schläfrigkeit“; (EFAS), Frohnhofen et al., 2010) were administered. Daily sleep diaries and actigraphy (GT3X, ActiGraph, Pensacola, FL, USA) were applied to monitor habitual bedtimes and wake times 7 days prior to the experimental nap sessions. These data verified that participants adhered to a regular sleep schedule.

### Study Design

The experimental procedure was identical to Ladenbauer et al. (2016). In brief, patients were tested in a balanced cross-over design in a stimulation and sham condition (n = 8 participants received so-tDCS and n = 8 received sham on the first experimental nap) that were separated by an interval of at least two weeks to prevent carry-over effects. Prior to experimental nap sessions, participants underwent an adaptation nap in the laboratory.

All nap sessions took place at the sleep laboratory of the Free University Berlin, Germany. Upon arriving at 11.30, participants were prepared for EEG-recordings, and then tested on two declarative memory tasks (a verbal paired-associate learning and visuo-spatial learning including picture and location memory) followed by a procedural memory task (finger sequence tapping). At 14.00, after a standardized small meal and preparation for so-tDCS, participants were asked to attempt to sleep during a period of 90 minutes, followed by memory tests thirty minutes after awakening.

Before learning and prior to retrieval after the nap, attentional capabilities (Test of Attentional Performance, TAP, Zimmermann and Fimm, 1995), the affective state (PANAS scales Watson et al., 1988), sleepiness (Tiredness Symptoms Scale, TSS, Bes et al., 1992 and Visual Analog Scale, VAS, Luria, 1975) and activation (VAS for tension) were assessed to control for possible confounding effects. For a schematic representation of the experimental procedure see Figure 1a.

### Slow oscillatory stimulation (so-tDCS)

The stimulation protocol was identical to that described previously by Ladenbauer et al. (2016). Stimulation electrodes (8 mm diameter) were positioned bilaterally at frontal locations F3 and F4 of the international 10-20 system (mounted into an EASY cap (Falk Minow Services, Munich, Germany)), with reference electrodes placed at each mastoid (ipsilateral; likewise 8-mm diameter). Anodal current was applied by a battery-driven stimulator (DC-Stimulator; NeuroConn, Ilmenau, Germany; current split to bilateral electrode sites) and oscillated sinusoidally at a frequency of 0.75 Hz (between 0 and 262.5 μA), resulting in a maximum current density of 0.522 mA/cm^2^. Electrode resistance was < 5 kΩ.

So-tDCS started four minutes after the subject had entered stable NREM sleep stage 2 and was delivered in a block-wise manner, each as a 5-min block of stimulation separated by stimulation free inter-block intervals of (at least) 1 min 40 s. The marker for the beginning of each 1-min stimulation-free interval (for analyses of immediate stimulation induced effects) was always set manually 40 s after the end of each stimulation period to exclude the strong and long lasting stimulation-induced drifts visible in our unfiltered online EEG signal from the analysis (interval of analysis will be referred to hereafter as “1-min stimulation-free interval”). As described in Ladenbauer et al. (2016), the number of the 5-min stimulation blocks (three to five blocks; three were required for inclusion) and the duration of the intermediate stimulation-free intervals depended on the individual subject’s sleep, as sleep was monitored following each stimulation block and each stimulation block was only initiated during NREM sleep stage 2, 3 or 4. As previous studies indicated that so-tDCS effects critically depend on ongoing brain state (Kanai et al., 2008; Kirov et al., 2009; Marshall et al., 2011), this protocol was chosen to account for higher sleep fragmentation in older adults to prevent application of so-tDCS during inappropriate brain states such as wake or sleep stage 1 phases.

During the sham session, stimulation electrodes were placed identical to the so-tDCS session, but the tDCS device remained off. The same criteria as for the so-tDCS condition were applied for the sham condition (first sham block began 4 min after onset of sleep stage 2; for subsequent sham blocks sleep stage 2 or slow wave sleep was required). Likewise the first 40 sec following each sham block were excluded from analyses. Participants were blinded for stimulation condition throughout the study. After completing all study-related procedures, they were asked whether they were able to guess in which experimental nap the stimulation had been applied and whether they felt any sensations during the naps.

### Sleep monitoring and preprocessing

During naps EEG was recorded from 26 scalp sites (FP1, FP2, AFz, F7, Fz, F8, FC5, FC1, FC2, FC6, C3, Cz, C4, T7, T8, CP5, CP1, CP2, CP6, P7, P3, Pz, P4, P8, O1, O2) using Ag-AgCl ring electrodes placed according to the extended 10-20 international EEG system. FCz was used as ground site. Data were recorded with the BrainAmp amplifier system (Brain Products GmbH, Munich, Germany) at a sample rate of 500 Hz and bandpass filtered between 0.05 and 127 Hz. All electrode recordings were referenced to an electrode attached to the nose. Impedances were less than 5 kΩ. Additionally, electromyograms at the chin as well as horizontal and vertical electrooculograms were recorded according to standard sleep monitoring.

Following the application of a notch filter (centered at 50 Hz with a bandwidth of 5 Hz), a combined semi-automated and visual rejection of raw data was applied to eliminate epochs contaminated by artifacts. This processing was done with BrainVision Analyzer software (Version 2.0, Brain Products, Munich, Germany).

### EEG analyses

We performed spectral as well as phase amplitude coupling analyses for the 1-min stimulation-free intervals following each so-tDCS and sham block. Depending on the number of actually performed stimulations or sham stimulations in a subject, three to five intervals were used for the analysis.

#### Spectral power

Spectral power was calculated for each 1-min stimulation-free interval per electrode using the Fast Fourier Transform on up to 11 overlapping (by 5 s) artifact-free segments each lasting 10 s. Corresponding intervals were used for the sham session. On each of these 10 s segments of EEG data, a Hanning window was applied before calculating the power spectra (frequency resolution 0.06 Hz). Subsequently, mean power (μν^2^) was calculated over the following frequency bands of interest: SO (0.5-1 Hz) and fast spindles (12-15 Hz). We additionally considered the slow spindle frequency band slow spindle (8-12 Hz) to compare with previous results (Marshall et al., 2006).

Topographic regions of interest (ROI) for these frequency bands were selected according to previous research (Mölle et al., 2011; Ladenbauer et al., 2016). The sites FC1, Fz, FC2 and CP1, Cz, CP2 were pooled into two ROIs, frontal and centro-parietal, respectively. Thus, mean spectral power reflects an average over the electrodes of each ROI. BrainVision Analyzer software (Version 2.0, Brain Products, Munich, Germany) was used to perform spectral power analyses.

#### Phase-amplitude coupling (PAC)

To assess the temporal relationships between memory-relevant oscillations in the EEG signal we employed an event-locked analysis based on Staresina et al. 2015, using the FieldTrip (Oostenveld et al., 2011) toolbox for MATLAB (The MathWorks, Natick, MA, USA) as well as custom MATLAB functions. Specifically, we characterized SO-to-spindle PAC for the frontal ROI (Fz, FC1, FC2) and the centro-parietal ROI (Cz, CP1, CP2) by applying the following procedure:

(i) SO events were identified for each subject, condition and electrode based on an established detection algorithm (Mölle et al., 2002, 2011). First, EEG data were filtered between 0.16-1.25 Hz (two-pass FIR bandpass filter, order = 3 cycles of the low cut-off frequency). Only artifact-free data were used and periods of wake state and REM sleep were also excluded for event detection. Second, periods of SO candidates were determined as time between two successive positive-to-negative zero-crossings in the filtered signal. For the next step events that met the SO duration criteria (period ≥ 0.8 s and ≤ 2 s, corresponding to 0.5–1.25 Hz) were selected. Third, event amplitudes were determined for the remaining SO candidates (trough-to-peak amplitude between two positive-to-negative zero crossing). Events that also met the SO amplitude criteria (≥ 75% percentile of SO candidate amplitudes, that is, the 25% of events with the largest amplitudes) were considered SOs. Finally, artifact-free epochs (−2.5 to +2.5 s) time-locked to the SO down-state in the filtered signal were extracted from the unfiltered raw signal for all events.

(ii) Time-frequency representations (TFRs) were calculated per event epoch and channel for frequencies 5-20 Hz in steps of 0.25 Hz using a sliding (10 ms steps) Hanning tapered window with a variable, frequency-dependent length (*mtmconvol* function of the FieldTrip toolbox). The window length always comprised a full number of five cycles to ensure reliable power estimates. TFRs were then normalized as difference to pre-event baseline (−2.5 to −1.2 s of the epoch) and averaged per subject, condition and ROI.

(iii) To quantify synchronization and locking between the (modulating) SO and the (modulated) oscillatory fluctuations of the fast spindle power we calculated the phase values of these time series and applied a synchronization index. Phase values were calculated for all time points of each extracted SO event and the corresponding fast spindle power fluctuation using the Hilbert transform. Spindle power fluctuation time series were obtained by the TFR bins averaged across the respective frequencies and up-sampled to the sampling frequency of 500 Hz. To ensure proper phase estimation, both SO and spindle power fluctuation time series were filtered beforehand in the range of the modulating SO event (0.5-1.25 Hz; two-pass FIR bandpass filter, order = 3 cycles of the low cut-off frequency). The synchronization index (SI) was then calculated between the two phase value time series for each event epoch and electrode. The resulting SI is a complex number of which the radius (r) indicates the strength of locking between the modulating SO and the modulated oscillatory fast spindle power fluctuation, and the angle (φ) represents the phase shift between these oscillations. In other words, φ indicates the phase of the SO at which fast spindle power is maximal across time. It was obtained by

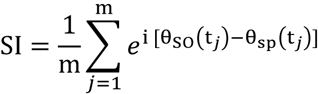

where m is the number of time points, θ_SO_(t_*j*_) is the phase value of the SO time series at time point t*j*, and θ_SP_(t_*j*_) is the phase value of the fluctuations of the fast spindle power time series at time point t*j* (Cohen, 2008). The interval for computing the SI was −1 s to +1 s around the SO center. The distribution of SIs per condition and ROI are visualized in Figure 3d. For statistical analyses (see below) SIs were averaged per subject, condition and ROI.

### Sleep architecture

The sleep structure including the time and proportion spent in different sleep stages was determined based on polysomnographic criteria according to Rechtschaffen and Kales (1968). For this purpose, EEG data was down-sampled to 250 Hz and 30-s epochs were scored manually by means of Schlafaus software (Steffen Gais, Lübeck, Germany) in sleep stages 1, 2, 3, 4 and REM sleep, epochs of wakefulness or movement artifacts. Epochs during so-tDCS were not scored due to strong artifacts from so-tDCS in the EEG signal. Likewise, corresponding epochs in the sham session were not scored, to obtain comparable time and proportions of sleep stages. Scoring for the 1-min stimulation-free intervals (between the 5-min blocks of acute stimulation) was additionally performed for 10-s epochs.

#### Memory tasks

All memory tasks were performed with Presentation software (Neurobehavioral Systems, Albany, CA, Version 14.8) and parallel versions were used for all tasks in the two experimental nap sessions. All memory tasks were also previously administered in Ladenbauer et al. (2016). In the visuo-spatial and verbal memory tasks subjects were instructed to memorize items for a later recall, but no specific strategy was recommended. There was no overlap regarding the stimuli used in the visuo-spatial and the verbal memory tasks.

### Visuo-spatial memory task

The visuo-spatial memory task consisted of 38 neutral pictures (objects, plants, scenes taken from the Affective Picture System (IAPS, Lang et al., 1998; MULTIMOST, Schneider et al., 2008) that appeared randomly at one of the four possible quadrants on the screen for 2 s with an inter-stimulus interval of 1 s (see Figure 1b). To account for primacy and recency effects four additional pictures (two at the beginning and two at the end) were included and disregarded in the analyses. Participants were instructed to memorize both the pictures (picture memory) and their locations (location memory). During recognition testing (prior to and after sleep), each picture (38 studied and 38 new pictures, random order) was displayed in the center of the screen (3 s), while participants indicated by button press whether they recognized each picture (“Old/New” decision). If participants recognized an item (“Old” decision), they were also required to indicate in which quadrant they believed the picture was presented during acquisition.

“Old/New” decisions in this task resulted in four possible response categories: hits (correct “Old” judgements), correct rejections (correct “New” judgements), false alarms (incorrect “Old” judgments) and misses (incorrect “New” judgements). As a measure for picture recognition memory accuracy percent correct responses was calculated as follows: proportion of hits + proportion of correct rejections. Further, potential response biases were considered by calculating the sum of the proportion of hits and false alarms.

For the accuracy of location memory, both correctly and incorrectly retrieved picture locations were taken into account: Number of correctly retrieved locations/number of hits – number of falsely retrieved locations/number of hits. Data from one participant was excluded from analyses for this task due to technical problems.

### Verbal memory task

Participants viewed 40 semantically related German word pairs (category-instance pairs: e.g. fruit-banana) that appeared centrally on the screen for 5 s with an inter-stimulus interval of 100 ms. Additional four word pairs were presented (two at the beginning and two at the end) to prevent primacy-recency effects. Following encoding, an initial cued-recall test was performed. The category name (cue) appeared centrally, and participants were instructed to say the respective stimulus (instance) word aloud after indicating (by button press within 10 s) whether they remembered the respective instance word. Subsequently the correct word pair was displayed for 2.5 s. This initial cued-recall test provided an additional learning opportunity, to help reach about 60% correct responses at the subsequent cued recall test. The immediate cued-recall and delayed cued-recall test (after sleep) were administered without additional presentation of the correct word pairs. Word pairs were presented in a different randomized order to prevent serial learning in each encoding and recall test. Participants’ verbal responses were recorded and cued recall performance was obtained by the proportion of the correct retrieved targets in the immediate and delayed recall test, respectively.

Moreover, we tested the impact of stimulation on interference rate (proportion of incorrect instance words in a nap session that were correct responses to the same categories in a previous session). Data from one participant was excluded from analyses for this task since she/he failed to follow task instructions.

### Procedural task

A sequential finger tapping task (SFTT; adapted from Walker et al., 2002) was used to investigate so-tDCS effects on procedural memory. Participants were asked to repeatedly tap a five-digit sequence (e.g. 4–2–3–1–4), that was displayed on the screen, with the non-dominant left hand as accurately and as quickly as possible within a 30-s interval (=trial). During pre-nap testing (learning), participants performed on twelve trails separated by 30-s breaks. Testing after nap (retrieval) contained four trials with breaks. Performance at learning and retrieval testing was determined by averaged correctly tapped sequences during the final three trials, respectively. Four participants failed to complete this task and were therefore excluded from analyses of this task.

### Statistical analyses

#### Spectral power

Differences between the two stimulation conditions (so-tDCS, sham) in the five outcome measures frontal and centro-parietal SO and fast spindle power as well as frontal slow spindle power were evaluated using a linear mixed model with random intercepts (LMM; Verbeke and Molenberghs 2000). This model was chosen to account for characteristic individual differences in sleep physiology (Buckelmüller et al., 2006; Tucker et al., 2007) and for unequal numbers of spectral power data points across subjects and condition (three to five applied so-tDCS and sham blocks depending on the individual subject’s sleep).These inequalities arised from the protocol chosen to prevent application of so-tDCS during inappropriate brain states.

In brief, the model is described by

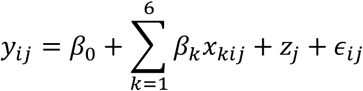

where *y_ij_* denotes the log-transformed spectral power value of participant *j* ∊{l,2,…,16} in interval *i ∊* {1,2,3,4,5 }, *β_0_* is the fixed intercept and *β_1_,…,β_6_* are the regression coefficients that correspond to the independent variables for *stimulation, x_1ij_* ∊ {0,1}, *baseline, x_2ij_* (see below), (centered) *time, x_3ij_* = *i -* 3, (centered) *time^2^, x_4ij_* = *(i -* 3)^2^, *time×stimulation, x_5ij_* = *(i-3)x_2ij_* and *time^2^×stimulation, x_6ij_* = *(i-3)^2^x_2ij_*. The subject specific random intercept is given by *zj* and *∊ij* denotes the residual (or random error). That is, the five time points (1-min stimulation-free interval following each stimulation block) were level-one units nested in subjects (level-two units). The model assumed that slopes were similar (no random slope model), since there was no previous evidence on inter-individual differences in stimulation effects (slopes) on sleep physiology. The *baseline* variable *(x_2ij_*) as covariate served to adjust for baseline differences in each frequency band, respectively. This variable took the subject-specific log-transformed spectral power value calculated from a 1-min interval preceding the first stimulation/sham block. As spectral power data did not exhibit a normal distribution they were log transformed prior to application of the LMM. Note that the values of the variables for *stimulation* and *baseline* did not vary across intervals. The squared centred *time* variable *(x_4ij_*) was incorporated to test for curvilinear course of SO and spindle power over intervals, as observed in healthy older adults (Ladenbauer et al., 2016). By the interaction *time×stimulation* we assessed whether the slopes of the curves differed between the stimulation conditions and an interaction term *time^2^×stimulation* was included to test whether the shape of the curves differed between the stimulation conditions.

### Phase-amplitude coupling

We tested for significant event-locked power changes in the TFR due to stimulation using a two-tailed paired-samples *t* test (group level statistics). To correct for multiple comparisons (−1.2 s to +1.2 s × 5-20 Hz) a cluster-based permutation procedure was applied as implemented in FieldTrip (Maris and Oostenveld, 2007; Staresina et al., 2015). The initial threshold for cluster definition was set to p < 0.025 and the final threshold for significance of the summed *t* value within clusters was set to p < 0.05.

To assess whether the (circular) SI angles were distributed non-uniformly with a specified mean we applied the V test (Berens, 2009). Using this test, the alternative hypothesis H1 states that the population is clustered around a known mean direction. In the current case we expected maximal fast spindle power around the SO up-state, that is, clustering around 0 deg (Staresina et al., 2015). Furthermore, we calculated the resultant vector length which indicates the degree of concentration of SI angles (Berens 2009). For a particular condition and ROI it is calculated by

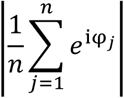

where *φ_j_* is the angle of the SI for subject *j* ∊ *{*1,…,*n}.*

### Sleep architecture

To evaluate so-tDCS effect on sleep structure t-tests (or Wilcoxon signed-ranks tests if indicated) were performed for the proportion of time spent in different sleep stages (in %) during the entire nap and during the 1-min stimulation-free intervals.

### Memory tasks

So-tDCS effects on retention performances in the memory tasks were tested by repeated measures analyses of variance (rmANOVA), including the within-subject factors STIMULATION (so-tDCS vs. sham stimulation) and TIMEPOINT (pre-nap vs. post-nap). The Greenhouse-Geisser correction of degrees of freedom was applied when appropriate.

Statistical analyses for memory tasks, power measures and sleep structure were performed using SPSS version 22.0 (SPSS, Chicago, IL, USA), while PAC behavior was statistically analyzed using the FieldTrip (Oostenveld et al., 2011) and CircStat (Berens, 2009) MATLAB toolboxes as well as custom MATLAB functions. A two-sided significance level α was set to 0.05 in all analyses. Given multiple testing for primary parameters of interest (picture memory, memory-related EEG spectral power, i.e., frontal and centro-parietal SO power and fast spindle power) we applied the Benjamini-Hochberg correction (Benjamini and Hochberg, 1995). To correct for multiple comparisons in the statistical TFR analysis, a cluster based permutation method was applied (see above). All other tests and comparisons were related to secondary hypotheses and p-values should be interpreted in a framework of exploratory analysis. Data are expressed as mean ± standard error of the mean (SEM), unless indicated otherwise.

## Author Contributions

N.K., A.F. and Ju.L. designed the study. Ju.L., R.d.B. and E.A. performed the experiments. Ju.L. and Jo.L. analyzed the data. Jo.L. contributed phase-amplitude coupling analyses. Ju.L., Jo.L. and U.G. performed statistical analyses. Ju.L., Jo.L. and A.F. wrote the paper.

## Acknowledgements

We would like to thank Sascha Tamm for technical support, and Sven Paßmann for helpful discussions. We further express our appreciation to the subjects who participated in the study. This work was supported by grants from the Deutsche Forschungsgemeinschaft (Fl 379-10/1, 379-11/1, SFB 910, DFG-EXC 257) and the Bundesministerium für Bildung und Forschung (FKZ 01EO0801, 01GQ1424A, 01GQ1420B).

